# Many erroneous noncoding transcripts in cancer cells can highly specifically regulate cancer-related genes and pathways

**DOI:** 10.1101/2024.07.13.603398

**Authors:** Sha He, Wei Xiong, Jianping Huo, Jie Lin, Jianmin Li, Hao Zhu

**Affiliations:** School of Basic Medical Sciences, Southern Medical University, Guangzhou, China; The First Affiliated Hospital, Nanchang University, Nanchang, China; College of Biological and Food Engineering, Guangdong University of Petrochemical Technology, Maoming, China; The Second Affiliated Hospital, Sun Yat-Sen University, Guangzhou, China

**Keywords:** Cancer, LncRNA, MSTRG transcripts, RNA-seq, Transcriptional regulation

## Abstract

Transcription and splicing errors in cancer cells generate erroneous transcripts. Since erroneous transcripts are degraded by the nonsense-mediated mRNA decay (NMD) pathway, whether they are junk or could be functional has been overlooked and understudied. We addressed this question by first performing a pan-cancer analysis and identified substantial erroneous noncoding transcripts (ENT) in cancers. Given that RNA/DNA binding domains (DBD) were predicted in ENTs, we deleted predicted DBDs in multiple ENTs in multiple cell lines, with RNA-sequencing and cell experiments before and after DBD deletion. DBD deletion caused significantly changed expression of ENTs’ target genes (whose promoter regions contain ENTs’ DNA binding sites, DBS) and changed cell migration and proliferation ability, indicating that many ENTs can transcriptionally regulate genes. Tightly coupled data analysis and experiments reveal that ENTs’ functions are highly cancer- and cellular-context specific, making ENTs a new class of safe and specific targets for noncoding RNA-based cancer therapeutics.

## 1. Introduction

RNA-sequencing (RNA-seq) has been widely used to study the cancer transcriptome for decades. Mature methods have been developed to map reads to genomes and assemble reads into transcripts. Nevertheless, since mammalian genomes are substantially transcribed, many assembled transcripts lack annotation. These transcripts, specifically labeled by assembling tools, may be normal transcripts in rare cell populations and specific time windows, or erroneous transcripts without functions, or erroneous transcripts with undesired functions. Since the nonsense-mediated mRNA decay (NMD) pathway degrades erroneous transcripts (with and without protein-coding ability) to protect cells from the consequences of transcription errors (Andjus et al., 2021; Lykke-Andersen and Jensen, 2015; Smith and Baker, 2015), and since most important genes and transcripts are assumed to have been annotated, unannotated transcripts (hereafter called erroneous transcripts) have been overlooked and understudied.

Transcription and splicing errors in cancer cells generate many erroneous transcripts (Bradley and Anczukow, 2023). Being largely understudied, whether they are harmless transcriptional noises or could have undesired functions remains an open question. Several recent studies revealed that some erroneous transcripts in cancer cells have functions (He et al., 2020; Liu et al., 2021; Yin et al., 2021). Since human lncRNA genes greatly outnumber protein-coding genes, many erroneous transcripts are noncoding sequences. Since RNA sequences function directly without being translated into proteins, erroneous noncoding transcripts (ENT) could be easier to obtain functions. Thus, examining ENTs in cancers is important for cancer biology. The question of whether ENTs have functions is important as well for considerable other diseases in which transcription and splicing errors widely occur.

To address this question, we first performed a pan-cancer analysis to re-analyze published raw RNA-seq data of cell lines and tissues of four cancer types. In this and following analyses, we used the *StringTie2* program (which uses ‘MSTRG’ to label unannotated transcripts) to assemble reads into transcripts, used the *Slnky* program to examine coding potential of MSTRG transcripts, used the *LongTarget* program to predict DNA-binding domains (DBD) in noncoding MSTRG transcripts and DNA-binding sites (DBS) in differentially expressed genes. This pan-cancer analysis revealed that many noncoding MSTRG transcripts (i.e., erroneous noncoding transcripts) have DBDs and DBSs and show correlated expression with target genes (genes with DBSs). We then deleted the 100-200 bp predicted DBD in three noncoding MSTRG transcripts in two cell lines, RNA-sequenced the wildtype (WT) and DBD knockout (KO) cell lines, and performed gene expression analysis. DBD knockout caused differential expression of many genes, especially predicted target genes of these noncoding MSTRG transcripts. To further validate, we conducted cell experiments to examine the migration and proliferation ability of the WT and KO cell lines. The KO cell lines showed significantly changed migration and proliferation ability compared with the WT cell lines. Finally, we developed a computational program to identify modules of regulated genes and related pathways in KO cell lines. The results revealed that modules of transcriptionally regulated genes are enriched for specific cancer-related pathways. Together, our results provide convincing evidence indicating that ENTs in cancer cells can specifically regulate gene expression. The cancer- and cellular context-specificity of this function makes ENTs a new class of safe and specific targets for noncoding RNA-based cancer therapeutics.

## 2. Results

### 2.1 Erroneous noncoding transcripts are generated pervasively in cancer cells

We downloaded the raw RNA-seq data of eight cancer cell lines (53 samples) and four cancer tissues (48 samples) with the corresponding controls from the GEO website (Supplementary Table 1). The cancer types include lung, liver, stomach, and colorectal cancer. We used the *StringTie2* program (which uses ‘MSTRG’ to label unannotated transcripts) to assemble reads into transcripts based on GRCh38 genome annotation (Pertea et al., 2015), used the *Slncky* program (which examines the coding potential of transcripts) to identify noncoding MSTRG transcripts (Chen et al., 2016), and detected differentially expressed genes and transcripts (logFC>1.0 and FDR< 0.05) (Supplementary Table 2). MSTRG transcripts were identified in all cancer cell lines and tissues, with 74% and 93% of MSTRG transcripts being cell-line and cancer tissue-specific (Fig. 1ABC).

**Figure 1.**
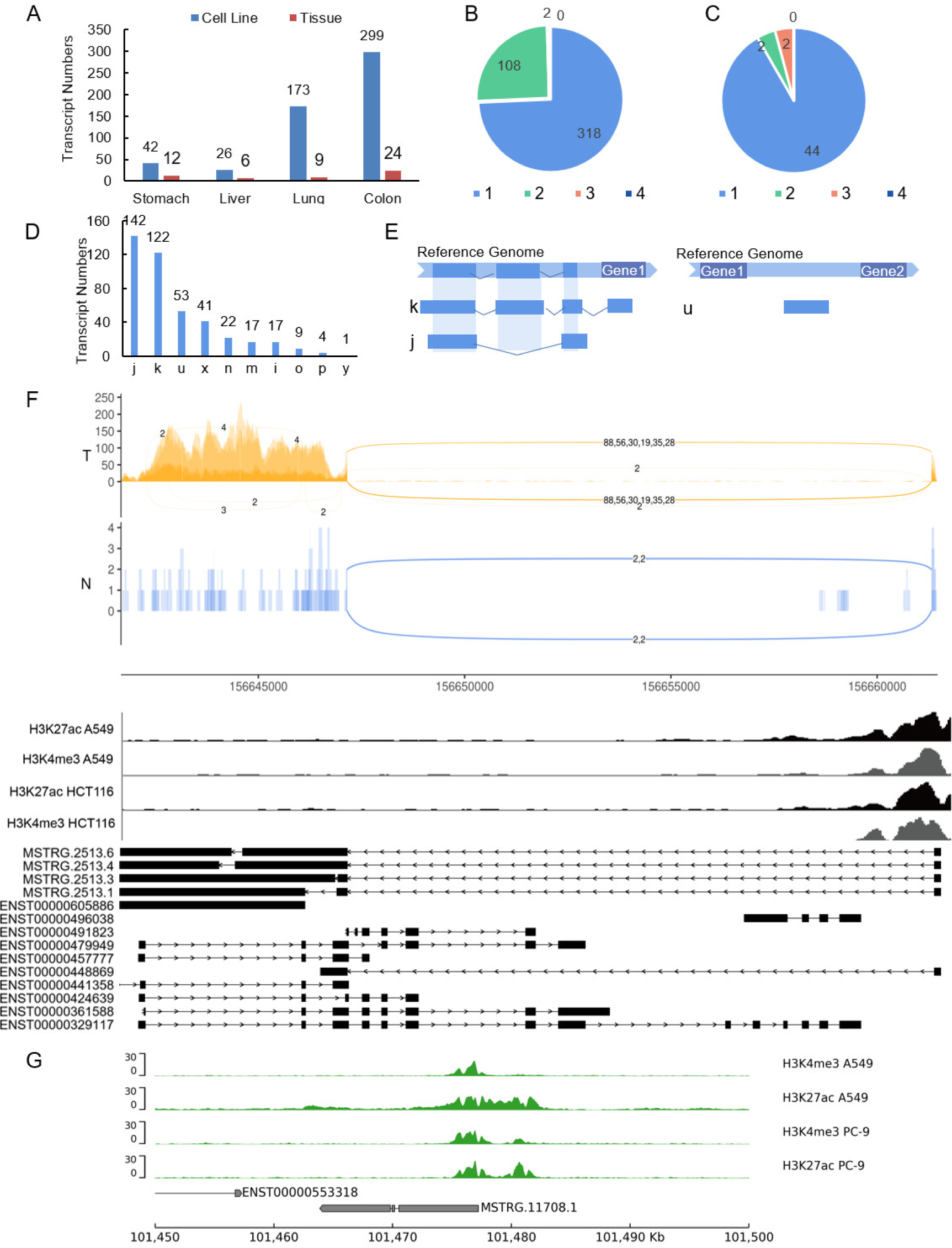
MSTRG transcripts are generated prevalently in cancer tissues and cell lines. (A) Numbers of MSTRG transcripts in the cell lines and tissues of the four cancer types. (B) Numbers of shared (indicated by numbers 2, 3, 4) and cancer-specific (indicated by number 1) MSTRG transcripts in the four cancer cell lines. (C) Numbers of shared and cancer-specific (indicated by numbers 1-4 as in panel B) MSTRG transcripts in the four cancer tissues. (D) Numbers of different types of MSTRG transcripts (see also Supplementary Fig. 2). (E) Illustrative exon distributions of the ‘j,’ ‘k,’ and ‘u’ type MSTRG transcripts. (F) From top to bottom are read coverage (‘T’ and ‘N’ indicate cancer and normal), H3K27ac and H3K4me3 signals, and the exon distribution of the MSTRG.2513 transcript family and the annotated transcripts. In the read coverage track, note the dense reads in cancer cells but sparse reads in normal cells in the region of the unannotated MSTRG.2513.1/3/4/6 exons. In the histone modification signal track, strong H3K4me3 and H3K27ac signals are at the first exon of MSTRG.2513. (G) Strong H3K4me3 and H3K27ac signals at the TSS of the ‘u’ type MSTRG.11708.1 in A549 and PC-9.

Erroneous transcripts can be generated by mechanisms including gene fusion, chimeric transcription, and alternative splicing. Especially, alternative splicing is present in 95% of multi-exon genes (Pan et al., 2008), 30% higher in cancers than in normal tissues (Kahles et al., 2018), and more common in lncRNAs than in mRNAs (Chen et al., 2021; Deveson et al., 2018). We found that many noncoding MSTRG transcripts overlap with annotated lncRNA genes (Supplementary Fig. 1). As examples, *CASC19*, a lncRNA gene up-regulated in many cancers (Ghafouri-Fard and Taheri, 2019; Wang et al., 2023), hosts multiple up-regulated noncoding MSTRG transcripts, and *FENDRR,* a lncRNA gene down-regulated in many cancers (Xu et al., 2014; Zheng et al., 2021), hosts multiple down-regulated noncoding MSTRG transcripts. These indicate that many noncoding MSTRG transcripts are products of wrong transcription and/or splicing of cancer-related lncRNA genes. We, therefore, used the *GffCompare* program to compare MSTRG transcripts with the related annotated genes and transcripts (i.e., references). *GffCompare* classifies MSTRG transcripts into 15 types (Supplementary Fig. 2) (Pertea and Pertea, 2020). The most prevalent ‘j’ and ‘k’ types overlap with annotated transcripts (Fig. 1DE), with 80% sharing transcription start site (TSS) and suggesting that many MSTRG transcripts are generated by wrong splicing. The ‘u’ type MSTRG transcripts in intergenic regions suggest their generation by mis-transcription. We measured the expression level of MSTRG transcripts using the percent spliced-in index (PSI) (Schafer et al., 2015), with PSI=1.0 and PSI<1.0 indicating constitutive exons (included in all transcripts) and reduced inclusion of alternative exons, respectively. Computed PSI values indicate that most MSTRG transcripts have higher expression levels than reference transcripts (Fig. 1F; Supplementary Fig. 3, 4), thus excluding the possibility that these MSTRG transcripts are assembly artifacts.

H3K4me3 and H3K27ac markers are often associated with activated promoters and enhancers and are widely used to identify transcriptional activation regions (Creyghton et al., 2010; Kimura, 2013). To determine the transcriptional activation regions of MSTRG transcripts, we collected and examined the H3K4me3 and H3K27ac histone modification data of the A549, HCT116, PC-9, and HpeG2 cell lines from the ENCODE website. Significant histone modification signals were found in 85% of ‘j’ and ‘k’ type MSTRG transcripts and 60% of ‘u’ type MSTRG transcripts at their TSS (Fig. 1FG; Supplementary Fig. 5, 6), indicating that wrong histone modification greatly promotes MSTRG transcript generation and further supporting that many MSTRG transcripts are not assembly artifacts.

### 2.2 DBDs and DBSs were predicted in noncoding MSTRG transcripts and differentially expressed genes, respectively

Many lncRNAs regulate transcription by binding to DNA sequences and recruiting epigenetic modification enzymes to binding sites. To examine whether noncoding MSTRG transcripts have DNA binding ability, we used the *LongTarget* program to predict DBDs in noncoding MSTRG transcripts and DBSs in promoter regions of differentially expressed genes (DEG) (Lin et al., 2019). DBDs were predicted in 157 of 172 up-regulated noncoding MSTRG transcripts in cancer cell lines, and 107 of these predicted DBDs were in unannotated exons. For example, three DBDs were predicted in unannotated exons in MSTRG.2513.6 (DBD1 is the DBD with most DBS), all having DBSs in many DEGs, and one DBD was predicted in an unannotated exon in MSTRG.4401.1, with DBSs in many DEGs (Fig. 2A-D). Genes and transcripts whose promoter regions contain a DBS of a MSTRG transcript are potential targets of the MSTRG transcript. The lengths of predicted DBDs (many > 100 bp) and the numbers of their DBSs (up to hundreds) suggest that these MSTRG transcripts have DNA-binding ability and may transcriptionally regulate many genes in proper contexts.

**Figure 2.**
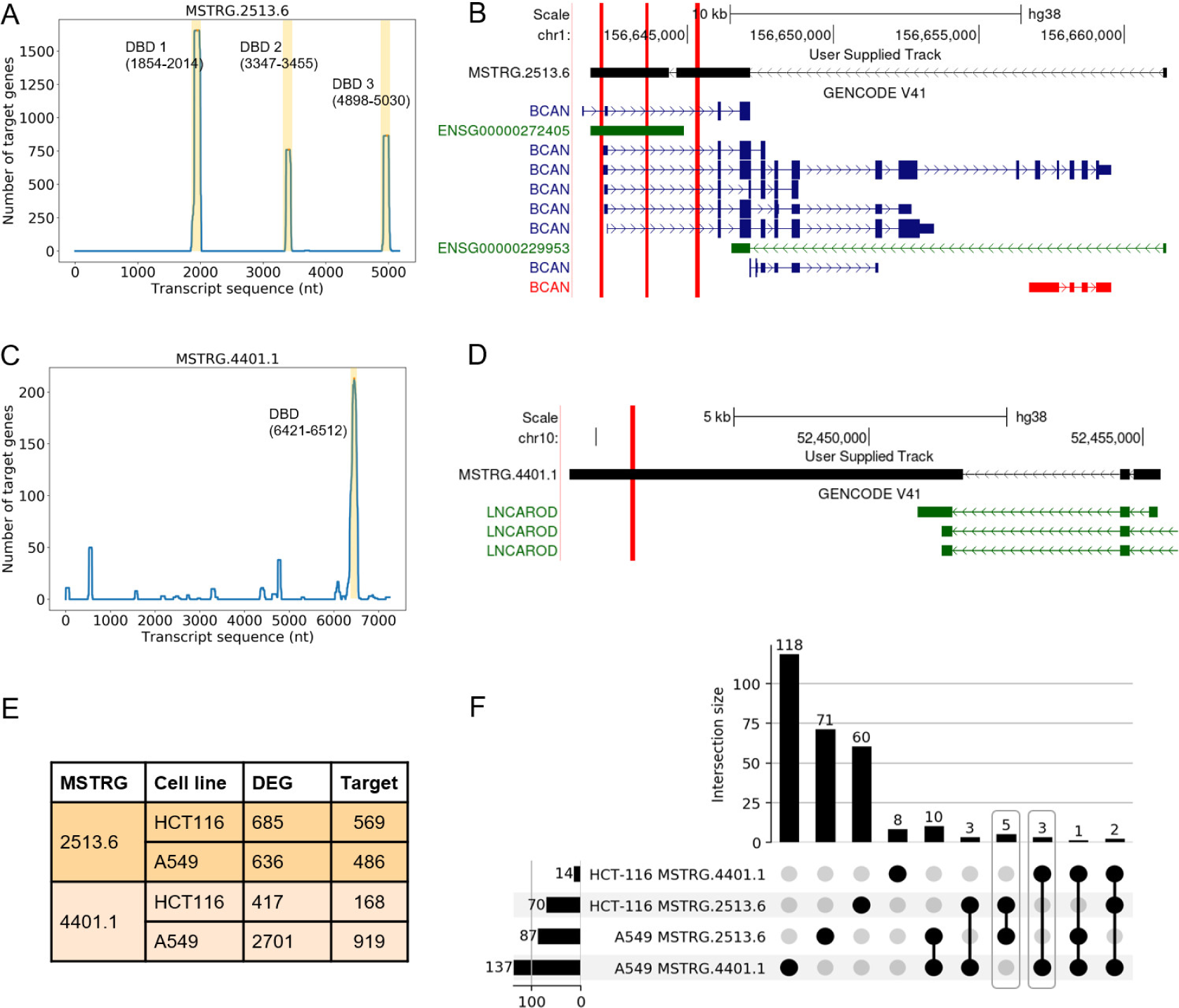
The DBD locations and DBS numbers of MSTRG.2513.6 and MSTRG.4401.1. (A) Three predicted DBDs (indicated by peaks) in MSTRG.2513.6 and the numbers of their DBS. (B) These DBDs (indicated by red bars) of MSTRG.2513.6 are in two unannotated exons. (C) The predicted DBD in MSTRG.4401.1 and the number of their DBS. (D) This DBD of MSTRG.4401.1 is in an unannotated exon. (E) Numbers of DEGs after MSTRG.2513.6 and MSTRG.4401.1 DBD knockout in A549 and HCT116; most DEGs are predicted targets of MSTRG.2513.6 and MSTRT.4401.1. (F) Numbers of the total (indicated by the left-side histogram), shared (indicated by the upper histogram), and specific (indicated by the dots) target genes in the 4 cases of DBD knockout.

### 2.3 DBD knockout caused significantly changed gene expression

To examine whether noncoding MSTRG transcripts use their DBDs to bind DNA sequences and regulate transcription, we used CRISPR/Cas9 to delete a 278 bp sequence containing DBD1 of MSTRG.2523.6 and a 170 bp sequence containing DBD1 of MSTRG.4401.1 in A549 and HCT116 (Supplementary Table 3). RNA-seq was performed before and after DBD knockout to analyze differential gene expression, and gel electrophoresis and read coverage analysis were made to validate DBD knockout (Supplementary Fig. 7, 8). After deleting DBD1 in MSTRG.2513.6 and in MSTRG.4401.1 in the two cell lines, hundreds to thousands of genes were differentially expressed compared with the WT cell lines. Notably, many of these DEGs were predicted target genes of MSTRG.2513.6 and MSTRG.4401.1; these DEGs were highly cancer and cellular context-specific (Fig. 2EF); |logFC| of target genes was significantly larger than |logFC| of non-target genes (Supplementary Fig. 9); and some MSTRG transcripts were completely activated or repressed by DBD knockout (Supplementary Table 4-8).

To examine what biological processes were influenced after MSTRG DBD knockout, we performed gene set enrichment analysis using multiple methods (e.g., *Metascape, gProfiler,* and *ClusterProfiler*) and databases (including the gene ontology (GO) database). All enrichment analyses generated similar results that are mutually supported (Supplementary Fig. 10, 11). For example, as *Metascape* reveals, in the cases of MSTRG.2513.6 DBD knockout in A549 and HCT116, DEGs were enriched in 19 gene sets, and the two most significantly enriched ones were “Extracellular matrix organization” (R-HSA-1474244) in the Reactome database and “cell-cell adhesion” (GO:0098609) in the GO database. In the cases of MSTRG.4401.1 DBD knockout in A549 and HCT116, DEGs were enriched in 18 gene sets, and the two most significantly enriched ones were “NABA CORE MATRISOME” (M5884) and “NABA BASEMENT MEMBRANES” (M5887) in the MSigDB database. Studies have revealed that the core extracellular matrix and structural components of the basement membrane are important for cancer development. The former provides structural and biochemical support to surrounding cells, therefore important for cell adhesion, migration, and proliferation; the latter is a thin and dense sheet of ECM, which is a structural barrier to cancer cell invasion (Chang and Chaudhuri, 2019; Socovich and Naba, 2019). Alterations in cell-cell and cell-ECM adhesion and in the organization and physical properties of the basement membrane make it permissive to cancer cell invasion (Glentis et al., 2017; Hamidi and Ivaska, 2018; Janiszewska et al., 2020).

### 2.4 DBD knockout changed cancer cells’ migration and proliferation ability

To further validate that noncoding MSTRG transcripts influence cancer cells’ phenotypes, we conducted cell experiments to examine whether the knockout of DBDs in MSTRG transcripts would change cancer cells’ migration and proliferation ability. First, the transwell migration assay revealed that the KO cell lines showed significantly stronger migration ability than the WT cell lines (Fig. 3A-D). Second, the cell proliferation assay revealed that the KO cell lines grew significantly faster than the WT cell lines (Fig. 3EG). Third, we used the cell apoptosis assay to detect and quantify cellular events associated with programmed cell death. We found that the KO cell lines showed increased cell apoptosis rates (Fig. 3FH). These results indicate that MSTRG.2513.6 and MSTRG.4401.1 influence multiple phenotypes of cancer cells.

**Figure 3.**
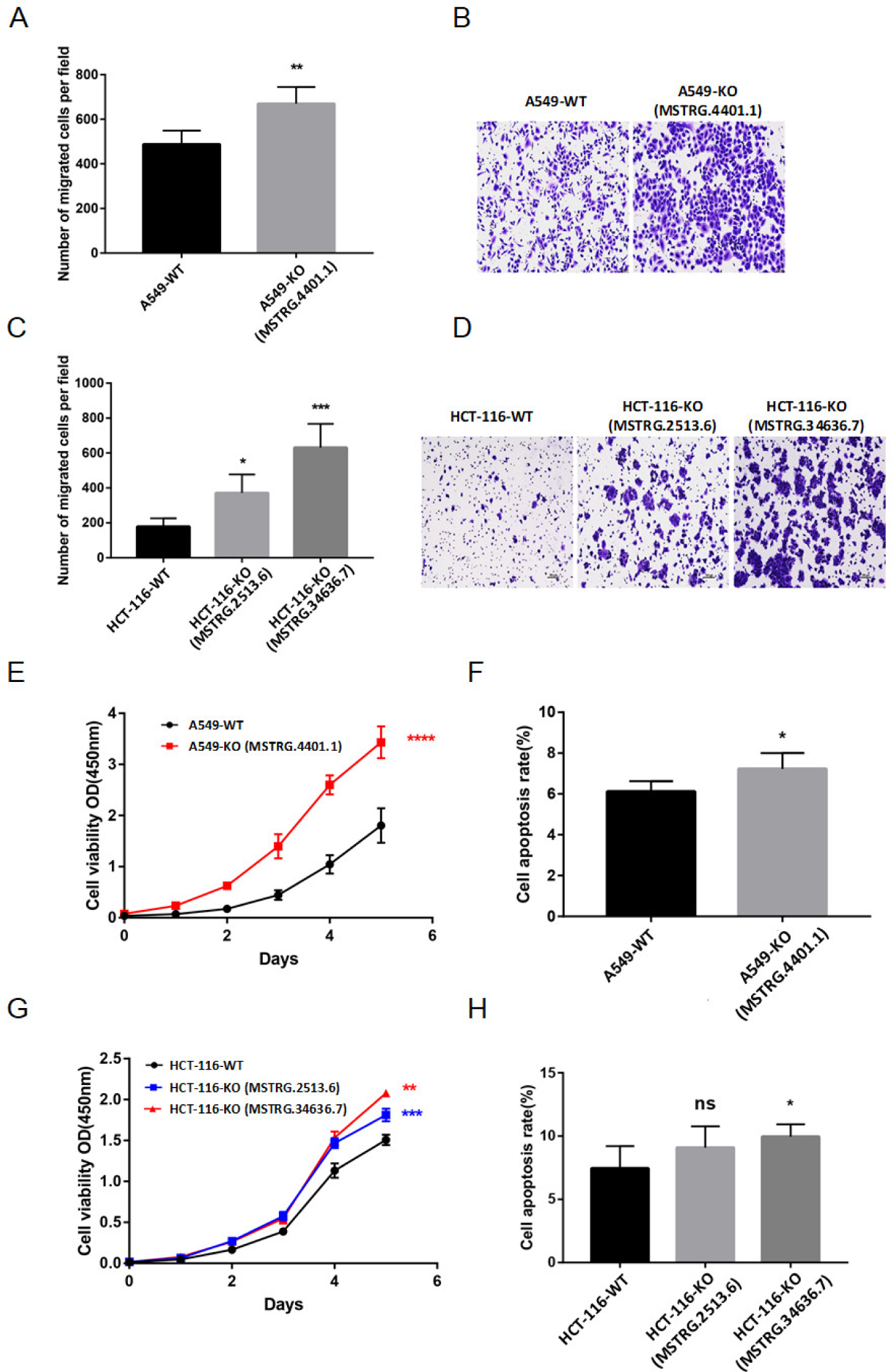
DBD knockout changed cancer cells’ migration, proliferation, and apoptosis properties. (AC) The transwell migration assay reveals that the KO cell lines obtain stronger migration ability (‘*’, ‘**’, and ‘***’ indicate P=0.012, 0.003, and 0.0001). (BD) The transwell migration assay reveals that the KO cell lines have a changed cell population. (EG) The cell proliferation assay using the “Cell Counting Kit-8” reveals that the two KO cell lines grew significantly faster (‘**’, ‘***’, and ‘****’ indicate P=0.008, 0.000124, and 0.000045). (FH) The cell apoptosis assay reveals that the KO cell lines showed a slightly increased cell apoptosis rate (‘ns’ indicates non-significant, ‘*’ indicates P=0.032).

### 2.5 Noncoding MSTRG transcripts function together with lncRNAs and specifically target genes in cancer-related pathways

Many studies use clustering methods to cluster genes with correlated expression into modules to reveal transcriptional regulatory mechanisms. Unsupervised clustering algorithms, however, cannot distinguish between regulators from targets (Saelens et al., 2018). To overcome this limitation, the *GRAM* program distinguishes regulatory TF sets and their target modules, but does not handle transcriptional regulation by lncRNAs (Bar-Joseph et al., 2003). To faithfully reveal transcriptional regulation, we developed a program (called *eGRAM*) that distinguishes regulatory lncRNA sets and their target modules and regulatory TF sets and their target modules based on combined targeting (TF/DBS and noncoding RNA/DBS) relationship and correlated expression relationship.

The inputs to *eGRAM* include differentially expressed genes, lncRNAs (including noncoding MSTRG transcripts) and predicted target genes, TFs and predicted target genes, and all KEGG and WikiPathways pathways. *eGRAM* revealed that noncoding MSTRG transcripts and lncRNAs form regulatory sets and regulate specific targets (parameters were Pearson coefficients>=0.8 and DBS >=100 bp). Target genes are highly cancer-, cellular context-, and MSTRG transcript-specific, thus showing enrichment in distinct cancer-related pathways (Fig. 4; Supplementary Fig. 12). In addition, in the four KO cases, many target gene modules are enriched in focal adhesion, PI3K-Akt signaling, and EGFR tyrosine kinase inhibitor resistance pathways, in line with the cell experiment and DEG analysis results. Some pathways (e.g., GnRH secretion, Porphyrin metabolism, and hematopoietic cell lineage pathways) have limited functional reports in cancer (Adapa et al., 2024; Limonta et al., 2012), suggesting that the impact of MSTRG transcripts on these pathways has not been fully investigated.

**Figure 4.**
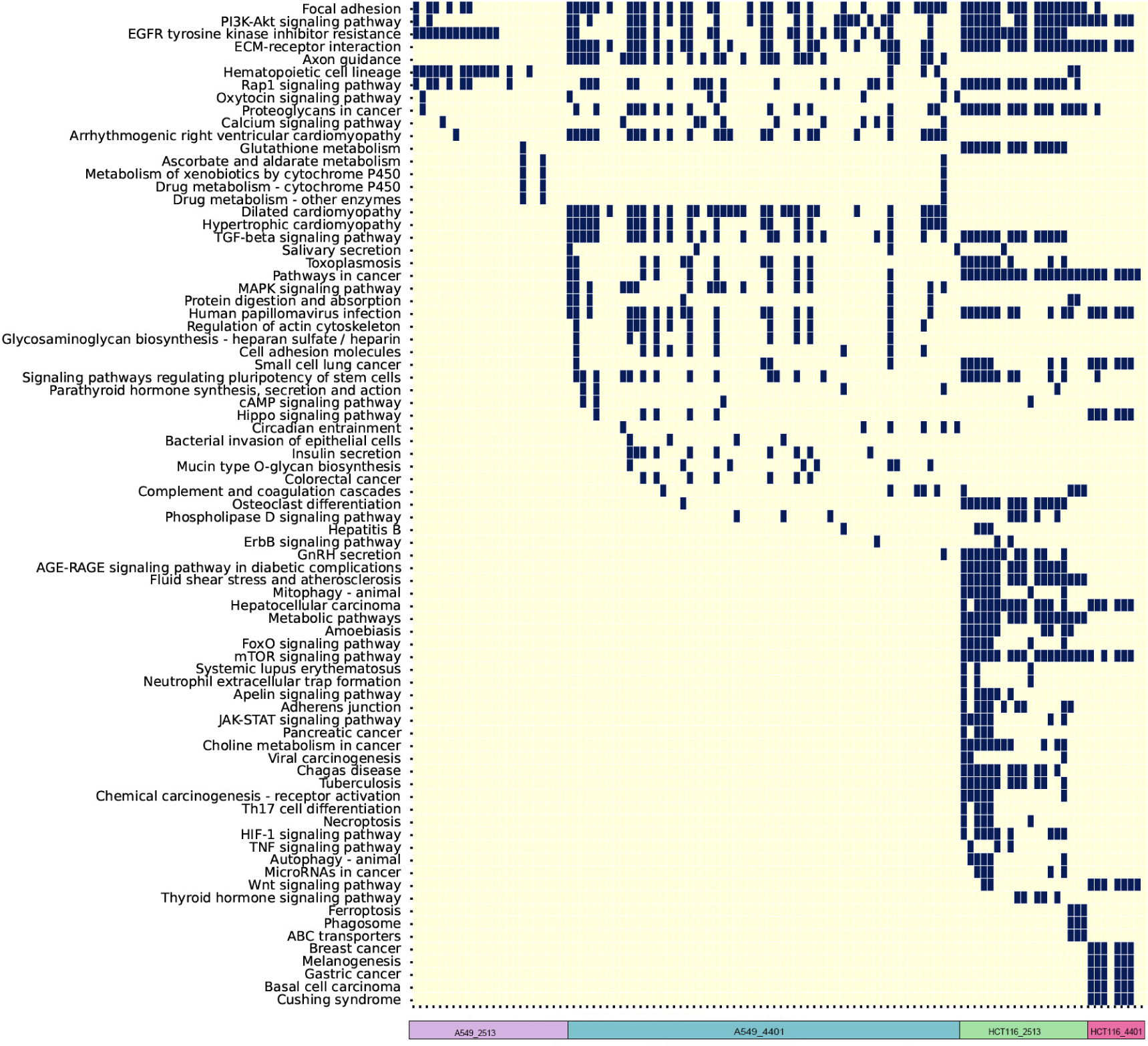
Target gene modules and enriched KEGG pathways in the four KO cases. (FDR<0.01). Left to right on the horizontal axis are modules in MSTRG.2513.6 KO A549, MSTRG.4401.1 KO A549, MSTRG.2513.6 KO HCT116, and MSTRG.4401.1 KO HCT116. Pathways enriched by only one module are not shown.

## 3. Discussion

Erroneous transcripts are prevalent in cancer cells (Bradley and Anczukow, 2023), some can escape from NMD (Lykke-Andersen and Jensen, 2015), and some may have functions (He et al., 2020; Li et al., 2019; Liu et al., 2021; Yin et al., 2021). These findings suggest that many erroneous transcripts, especially noncoding ones, are functional rather than junk, and this may be an important and understudied aspect of cancer biology. From a pan-cancer analysis to deletion of DBDs in noncoding MSTRG transcripts, this study combined computational and experimental approaches and systematically examined erroneous noncoding transcripts in multiple cancers. The results clearly indicate that erroneous noncoding transcripts may function like lncRNAs and specifically regulate cancer-related genes. This finding may be of great interest, as it reveals an important but overlooked aspect of cancer transcriptome and possibly the transcriptomes of many other diseases that are also characterized by transcriptional and splicing errors. This finding also helps explain the poor reproducibility of many cancer studies, as the highly cellular context-specific functions of MSTRG transcripts have been overlooked. The high cancer and cellular context specificity of MSTRG transcripts and their functions suggest they can be a class of new and safe diagnostic and therapeutic targets, as targeting them can avoid the adverse effects of targeting important proteins or mRNAs (Jing et al., 2022). Therapeutic targeting of noncoding RNAs represents an attractive approach for the treatment of cancers as well as many other diseases (Winkle et al., 2021).

MSTRG transcripts have attracted researchers’ attention and have been examined recently. Compared with knocking down MSTRG transcripts using RNA interference (Li et al., 2019; Liu et al., 2021; Yin et al., 2021), deleting the 100-200 bp predicted DBDs is more accurate for revealing noncoding transcripts’ transcriptional regulatory functions. In this study, the coverage of RNA-seq reads, the significantly changed target gene expression, and the changed cell migration and proliferation ability firmly validate and support the functions of DBDs and, thus, noncoding MSTRG transcripts.

This study also raises several questions to be further explored. First, considerable erroneous noncoding transcripts may be true transcriptional noise; distinguishing them from those that are functional is a question; and predicting whether they contain a strong DBD is a strategy. Second, whether noncoding MSTRG transcripts are mainly oncogenic or tumor suppressor remains unclear. Third, whether functional MSTRG transcripts are also generated in cells of other diseases, especially neurodegenerative diseases.

## 4 Materials and methods

### 4.1 Reagents and resources

|  |  |
| --- | --- |
| RPMI 1640 media | BasalMedia L210KJ<br>(www.basalmedia.com) |
| DMEM media | BasalMedia L110KJ<br>(www.basalmedia.com) |
| FBS (Fetal bovine serum) | GIBCO 10270-106 (www.thermofisher.cn) |
| Penicillin-Streptomycin | BasalMedia S110JV<br>(www.basalmedia.com) |
| 0.25% Trypsin-EDTA solution | BasalMedia S310KJ<br>(www.basalmedia.com) |
| Transwell chamber | Falcon 353097 (ecatalog.corning.com) |
| PBS (Phosphate-buffered saline)<br>buffer | ZSGB-BIO ZLI-9062 (www.zsbio.com) |
| 4% paraformaldehyde | Biosharp BL539A (www.biosharp.cn) |
| Crystal violet staining solution | Beyotime C0121 (www.beyotime.com) |
| Cell Counting Kit-9 | Dojindo CK04 (www.dojindo.cn) |
| Annexin V APC Apoptosis Detection<br>Kit | BioGems 62700-80-100 (www.bio-gems.com) |
| 10-cm Cell culture dish | Jetbiofil TCD010100 (www.jetbiofil.com) |
| 24-well Cell culture plate | Jetbiofil TCP010024 (www.jetbiofil.com) |
| 6-well Cell culture plate | Jetbiofil TCP010006 (www.jetbiofil.com) |
| 96-well Cell culture plate | Jetbiofil TCP010096 (www.jetbiofil.com) |

### 4.2 Data collection

The RNA-seq data of four cancer types (i.e., lung, liver, stomach, and colorectal cancer) were downloaded from the GEO website (https://www.ncbi.nlm.nih.gov/geo/). Data include cancer cell lines and the corresponding normal cell lines, and cancer tissues and the corresponding normal tissues (Supplementary Table 1). The cancer and the corresponding normal tissues were from 24 patients (cancerous tissues and para-cancerous tissues). Gender, age, and tumor stage were neglected. The ChIP-seq data (H3K4me3 and H3K27ac, bigwig file) for A549, HCT116, HpeG2, and PC-9 cell lines were downloaded from the ENCODE portal with the following identifiers: ENCSR636PIN, ENCSR778NQS, ENCSR000EUS, ENCSR000EUT, ENCFF084DIM, ENCFF456IJK, ENCFF556HVB, ENCFF700JHP (https://www.encodeproject.org/) (Luo et al., 2020).

### 4.3 Assembling reads and identifying noncoding MSTRG transcripts

First, we used the *Trim_Galore* program (with default paraments) to trim adaptor sequences and screen low-complexity or low-quality sequences. Second, we used the *HISAT2* program (with default paraments) to align clean sequencing reads to the human genome GRCh38 (Kim et al., 2019). Third, we used the *Stringtie2* program (with default paraments) to assemble aligned reads into transcripts based on GRCh38 genome annotation (Kovaka et al., 2019). *StringTie2* uses ‘MSTRG’ to label unannotated transcripts. Fourth, we merged (using the ‘Transcript merge mode’ of *StringTie2*) assembled transcripts in all cell line samples and all tissue samples, respectively, into a set of non-redundant transcripts (which makes MSTRG transcripts at the same coordinates in all cell line and tissue samples consistently labeled). Fifth, we estimated expression levels of assembled transcripts and used the *prepDE.py* program (with default parameters) to extract read count information directly from the files. Transcripts in less than 3 samples with CPM<1 were considered noise and filtered. Sixth, since batch effects caused by biological experiment conditions may influence statistical power and induce spurious differences between study groups (Nygaard et al., 2016), we used the *Combat_seq* program from the *sva* R package to remove the batch effects between samples in the same group (Leek et al., 2012). Seventh, we used the *Slncky* program (with ‘--no_collapse, --no_self’ paraments) to examine the coding potential and identify noncoding MSTRG transcripts. *StringTie2* and *Slncky* classified transcripts into four classes: annotated protein-coding, unannotated protein-coding, annotated noncoding, and unannotated noncoding.

### 4.4 Analyzing and visualizing alternative splicing and transcript abundance

We used the *SUPPA2* program (with default parameters) to calculate the isoform percent spliced-in (PSI) for each noncoding MSTRG transcript based on TPM (Trincado et al., 2018). The PSI was calculated as PSI = IR/(IR + ER), with IR and ER being inclusion and exclusion reads (Schafer et al., 2015). We used the *Ggsashimi* program (with default parameters) to visualize all alternative splicing in transcripts at a given locus (Garrido-Martin et al., 2018). We used the *Corrplot* program (with default parameters) to plot the percent of the target genes in DEGs (https://github.com/taiyun/corrplot). To draw the histone modification signal map, we first used the *DeepTools ComputeMatrix* program to calculate scores per genome region from the Bigwig file (Ramirez et al., 2016), by which we prepared an intermediate file to be used with the *plotProfile* program. Then, we used the *plotProfile* program to create a plot for scores over sets of genomic regions provided by the BED file. The genomic region was set to 5 Kbps upstream and downstream of the transcription start site.

### 4.5 Classifying MSTRG transcripts

We used the *GFFcompare* program (with default parameters) to classify and annotate transcripts according to their location to the closest reference transcripts (Pertea and Pertea, 2020). Specifically, ‘u’ indicates intergenic unknown transcripts, and ‘j’, ‘k’, ‘m’, and ‘n’ indicate MSTRG transcripts generated by alternative splicing.

### 4.6 Predicting MSTRG transcripts’ DBDs, DBSs, and target genes and predicting TFs’ DBSs

*LongTarget* can more robustly and accurately identify DBS than other methods (Wen et al., 2022). We used the *LongTarget* program (with default parameters) to predict DNA-binding domains (DBDs) in noncoding MSTRG transcripts and DNA-binding sites (DBSs) in the promoter regions (1500 bp upstream and downstream of the transcription start site) of differentially expressed genes (Lin et al., 2019). A gene that is differentially expressed in normal and cancer cells, whose promoter region contains the DBS of a noncoding MSTRG transcript, and whose expression is highly correlated with the expression of the noncoding MSTRG, was assumed to be a target gene of the noncoding MSTRG transcript. We used the *CellOracle* program (with default parameters) to predict DBSs of differentially expressed TFs (Kamimoto et al., 2023; Lin et al., 2019).

### 4.7 Knocking out sequences containing DBDs

A revised version of CRISPR/Cas9 (CRISPR-U^TM^) was used to knock out a short DNA sequence that contains the predicted DBD in noncoding MSTRG transcripts. Six cases of knockout were performed by BUIGENE, Guangzhou, China (http://www.ubigene.com). The following criteria and steps were taken to design the gRNA sequences. First, we used the *CRISPOR* program (with default parameters) to detect candidate target sites and evaluate the specificity scores (Haeussler et al., 2016); the sites with scores greater than 60 were selected. Second, we used the *CCTop* program (with default parameters) to assess the sgRNA efficacy score (Labuhn et al., 2018; Stemmer et al., 2015); the sites with efficacy scores exceeding 0.56 were selected (Supplementary Table 3). Sequence knockout was validated experimentally. First, PCR amplification for targeted regions and agarose gel electrophoresis were performed to confirm the knockout results (Supplementary Fig. 7). Second, RNA-seq was performed before and after the knockout; the knockout regions show low reads coverage (Supplementary Fig. 8).

### 4.8 RNA-sequencing of cell lines and identifying differentially expressed genes

Before and after DBD knockout, RNA-sequencing (RNA-seq) was performed to detect gene expression (the MSTRG.34636.7 KO HCT cell line did not perform RNA-seq, see Fig. 3). Total RNA was extracted from each sample, and rRNAs were removed. RNA sequencing was performed by HaploX, Shenzhen, China (https://www.haplox.cn/).

The *EdgeR* program was used to determine and analyze differentially expressed transcripts and genes (DETs and DEGs) between WT and KO cell lines and between normal and cancer tissues (Robinson et al., 2010). DETs and DEGs were identified upon |logFC|>1 and FDR<0.05.

### 4.9 Analyzing functional enrichment of differentially expressed target genes

We used multiple programs, including *Metascape*, *gProfiler*, and *ClusterProfiler4.0* and multiple databases, including Gene Ontology (GO), KEGG, WikiPathways, Reactome, and Molecular Signatures Database (MSigDB), to perform functional enrichment of differentially expressed target genes. We also performed gene set and pathway enrichment analysis for each gene module detected by the *eGRAM* program (Fig. 4). The *eGRAM* program currently contains 353 KEGG pathways and 899 WikiPathways pathways.

### 4.10 Identifying the regulators/targets relationship

Transcriptional regulators regulate target genes with specific mechanisms (e.g., TF-TF’s DBS, lncRNA-lncRNAs’ DBS). As an upregulated regulator up/down-regulates its targets, its targets’ expression is highly correlated with the regulators’ expression. Thus, gene expression analysis should explore both correlation and regulation (e.g., targeting) relationships. Clustering algorithms (e.g., *WGCNA*) identify modules of genes upon correlation without considering targeting relationships (Saelens et al., 2018). The *GRAM* program identifies modules of genes upon both correlation and the TF-target relationships but does not handle lncRNAs (Bar-Joseph et al., 2003). We developed the *eGRAM* program that explores correlation and the two kinds of targeting relationships (TFs and their targets upon TF-DNA binding and lncRNAs and their targets upon lncRNA-DNA binding). *eGRAM* identifies regulatory lncRNA sets upon correlation (between lncRNAs, including noncoding MSTRG transcripts), regulatory TF sets upon correlation (between TFs), and modules of targets upon correlation and targeting relationships (both between regulators and targets). The inputs to *eGRAM* include gene expression values, lncRNA-target relations, TF-target relations, and all pathways in the KEGG and WikiPathways databases. These pathways enable the identification of target modules’ biological functions. The identified modules should be theoretically reliable, especially when related parameters are high. In our analysis, the cutoffs for lncRNA DBS and TF DBS were 100 and 10, respectively; the cutoff for Pearson correlation was 0.8 between regulators (lncRNAs, TFs) and between regulators and targets (lncRNAs/TFs and targets); module size cutoff was 50; and FDR was 0.01.

*eGRAM* performs the following steps. (a) Accept three input files (gene expression values, lncRNAs’ DBSs at potential target genes, and TFs’ DBSs at potential target genes) (Supplementary Table 4-7, 9-16) and parameters. (b) Identify each lncRNA’s (noncoding MSTRG transcripts were treated as lncRNAs) correlated lncRNAs, which form a regulatory lncRNA set. (c) Identify each TF’s correlated TFs, which form a regulatory TF set. (d) Compute the correlation between each lncRNA and all genes and between each TF and all genes. (e) Identify each lncRNA’s target genes and each TF’s target genes. (f) Identify each regulatory lncRNA set’s target module upon correlation and targeting relationship, and identify each regulatory TF set’s target module upon correlation and targeting relationship. (g) Collect modules with size>50. (h) Check whether TFs’ modules contain lncRNAs’ targets and whether lncRNAs’ modules contain TFs’ targets, revealing whether genes and modules are co-regulated by lncRNAs and TFs. (i) Perform pathway enrichment analysis for each module using hypergeometric distribution test and all (353) KEGG pathways and (899) WikiPathways pathways.

### 4.11 Cell experiments protocols

#### Transwell migration assay

First, digest cells in the culture dish with trypsin, terminate the digestion with a complete culture medium, centrifuge cells at 1000 rpm for 5 minutes, resuspend cells in the culture medium, and count cells. Second, take a new 24-well plate, add 800µl of culture medium containing 20% FBS, place the Matrigel-coated Transwell chamber in a 24-well plate containing the culture medium, evenly seed 200µl of culture medium containing 1×10^5 cells in the upper chamber of the Transwell, and culture in an incubator, avoiding the formation of bubbles throughout the process. Third, the culture medium for cell lines HCT116-WT and HCT116-2513-KO is RPMI 1640, with a culture time of 48 hours; the culture medium for A549-WT and A549-4401-KO is DMEM, with a culture time of 33 hours. Fourth, after the culture is complete, remove the chamber using a tweezer, discard the culture medium, and gently wash the chamber three times with PBS. Fifth, add 800µl of 4% paraformaldehyde to new wells in a new 24-well plate to fix the cells for 15 minutes. After fixation, transfer the chambers to new wells containing 500µl of 0.1% crystal violet staining solution prepared with pure water for staining for 15 minutes. After staining, rinse the chamber with pure water and gently wipe the cells and moisture from the upper chamber with a cotton swab. Sixth, after the chamber air-dry naturally, place it under an upright optical microscope for observation and photography. Two replicates are made.

#### Cell proliferation assay using Cell Counting Kit-8

First, cells are digested with trypsin, centrifuged at 1000 rpm for 5 minutes, resuspended in culture medium, and counted. Second, 200 µl of culture medium containing 1×10^3 cells is evenly added to each well of a 96-well plate, with 5 replicate wells per cell group. To prevent liquid evaporation within the wells, a suitable amount of sterile PBS is added around the periphery of the 96-well plate. The plate is placed in an incubator. After the cells adhere to the well surface, the original culture medium in each well is discarded. Subsequently, 100 µl of culture medium containing 10% CCK-8 is added to the 5 wells of each cell strain, avoiding bubbles throughout. Third, a blank control group is set up simultaneously to minimize background errors caused by the detection reagent. Fourth, after a light-protected incubation for 2 hours, each well’s absorbance at 450nm is measured using a microplate reader. The absorbance value at day 0 is recorded as the initial value, and absorbance values are continuously measured for 6 days, from which a proliferation curve is plotted using the absorbance values.

#### Cell apoptosis assay

First, wash the trypsin-digested cells with PBS and centrifuge at 1000rpm for 5 minutes. Second, resuspend the cells in Annexin V Binding Buffer to a concentration of 1×10^6/ml, take 100 µl of the cell suspension into a new EP tube, and add 5 µl of Annexin V and 7-AAD staining solution to each sample, followed by light-protected incubation at room temperature for 15 minutes. Third, after incubation, add 400 µl of Annexin V Binding Buffer for resuspending the cells and detect cells using flow cytometry within 1 hour. Fourth, count and compare the proportion of Annexin V APC+ cells in each cell group. Five replicates were made.

## AUTHOR CONTRIBUTIONS

H.Z. conceived the study and drafted the manuscript. S.H., W.X., H.Z. analyzed data. J.H. and J.L. performed cell experiments. J.L. and H.Z. developed the *eGRAM* program. J.L. and H.Z. supervised the study and revised the manuscript. All authors have read the manuscript and consent to its publication.

## ACKNOWLEDGEMENTS

This work was supported by the National Natural Science Foundation of China (31771456).

## DATA AND CODE AVAILABILITY

The gene expression data downloaded from the GEO website are publicly available. The raw RNA-seq data of the four DBD knockout cases have been deposited in the NCBI GEO database (https://www.ncbi.nlm.nih.gov/geo) (the accession number is GSE229846). The *LongTarget* program is available at (http://www.gaemons.net/). The *StringTie2, Slnky,* and *CellOracle* programs are publicly available (see related citations). The *eGRAM* program is available at the GitHub website (https://github.com/LinjieCodes/eGRAMv2R1). Other data are in Supplementary Tables.

## SUPPLEMENTARY DATA

Two supplementary files—one PDF file containing 10 supplementary figures and one Excel file containing 16 supplementary tables — are available with the manuscript online.

