## Supplementary Figures for "Many erroneous noncoding transcripts in cancer cells can highly specifically regulate cancer-related genes and pathways"

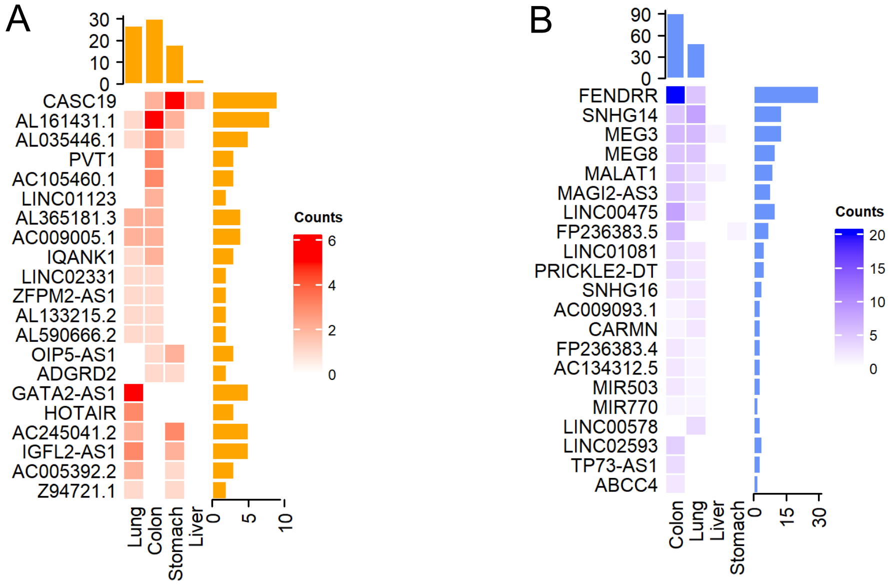


Supplementary Figure 1. The top 20 lncRNA genes that host noncoding MSTRG transcripts in different cancers. The top yellow histogram indicates the number of noncoding MSTRG transcripts in the four cancers; the right-side yellow histogram indicates the number of noncoding MSTRG transcripts in lncRNA genes; the rightmost color bar indicates the number of noncoding MSTRG transcripts in each lncRNA gene in each cancer. (A) The top 20 lncRNA genes that host up-regulated noncoding MSTRG transcripts. (B) The top 20 lncRNA genes that host down-regulated noncoding MSTRG transcripts.

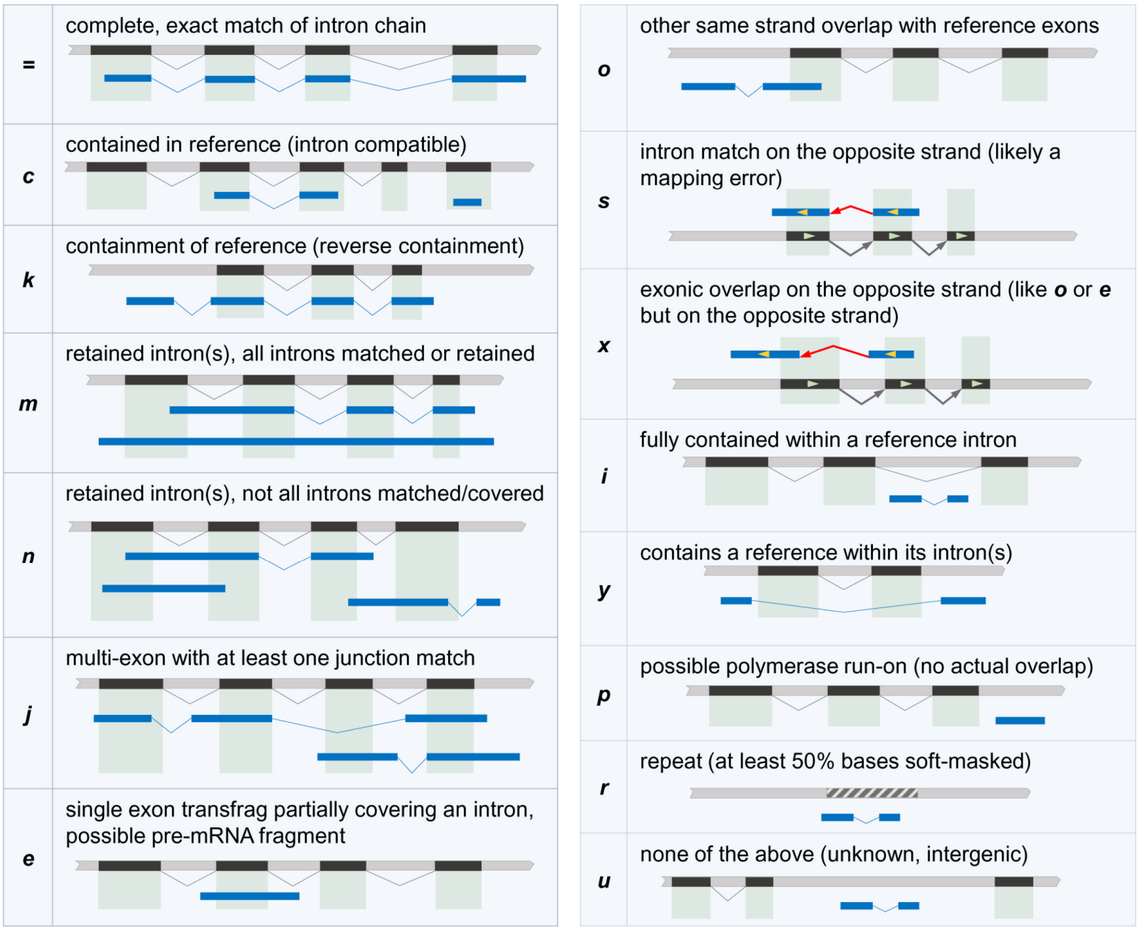


Supplementary Figure 2. Types of noncoding MSTRG transcripts (from Pertea and Pertea (2020) with permission).

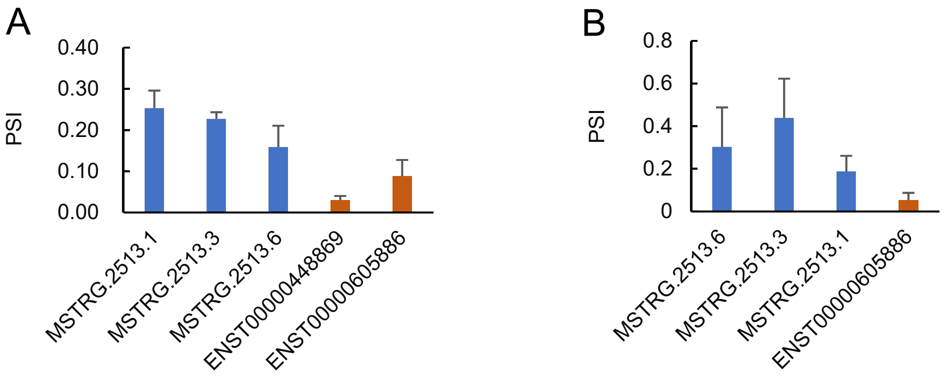


Supplementary Figure 3. The expression levels of MSTRG.2513 members as well as the overlapping annotated transcripts. (A) In the HCT116 cell line. (B) In the A549 cell line.

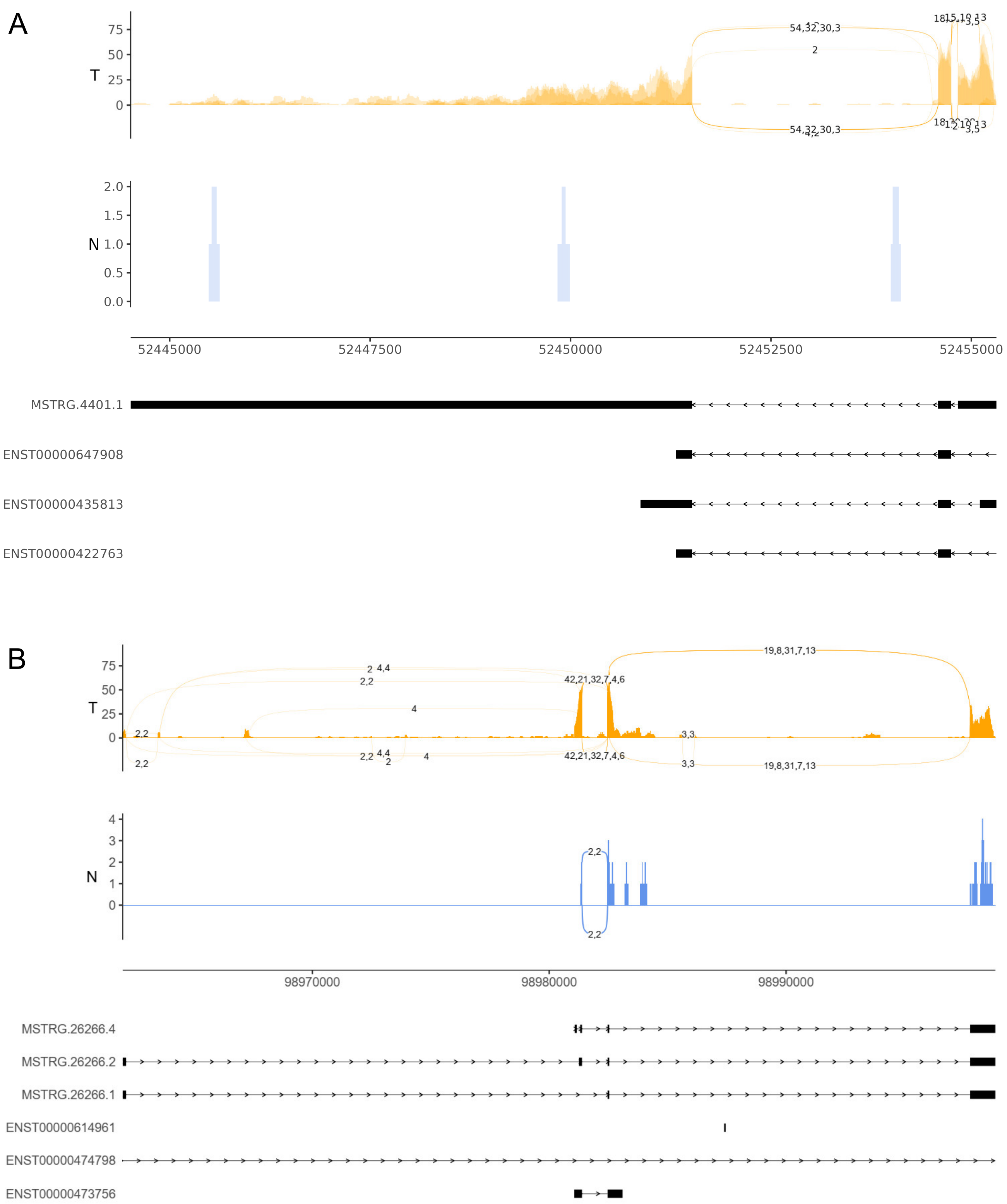


Supplementary figure 4. Reads coverage and exon distribution of MSTRG transcripts at annotated transcripts. 'T' and 'N' indicate cancer and normal cell lines. (A) MSTRG.4401.1. (B) MSTRG.26266.

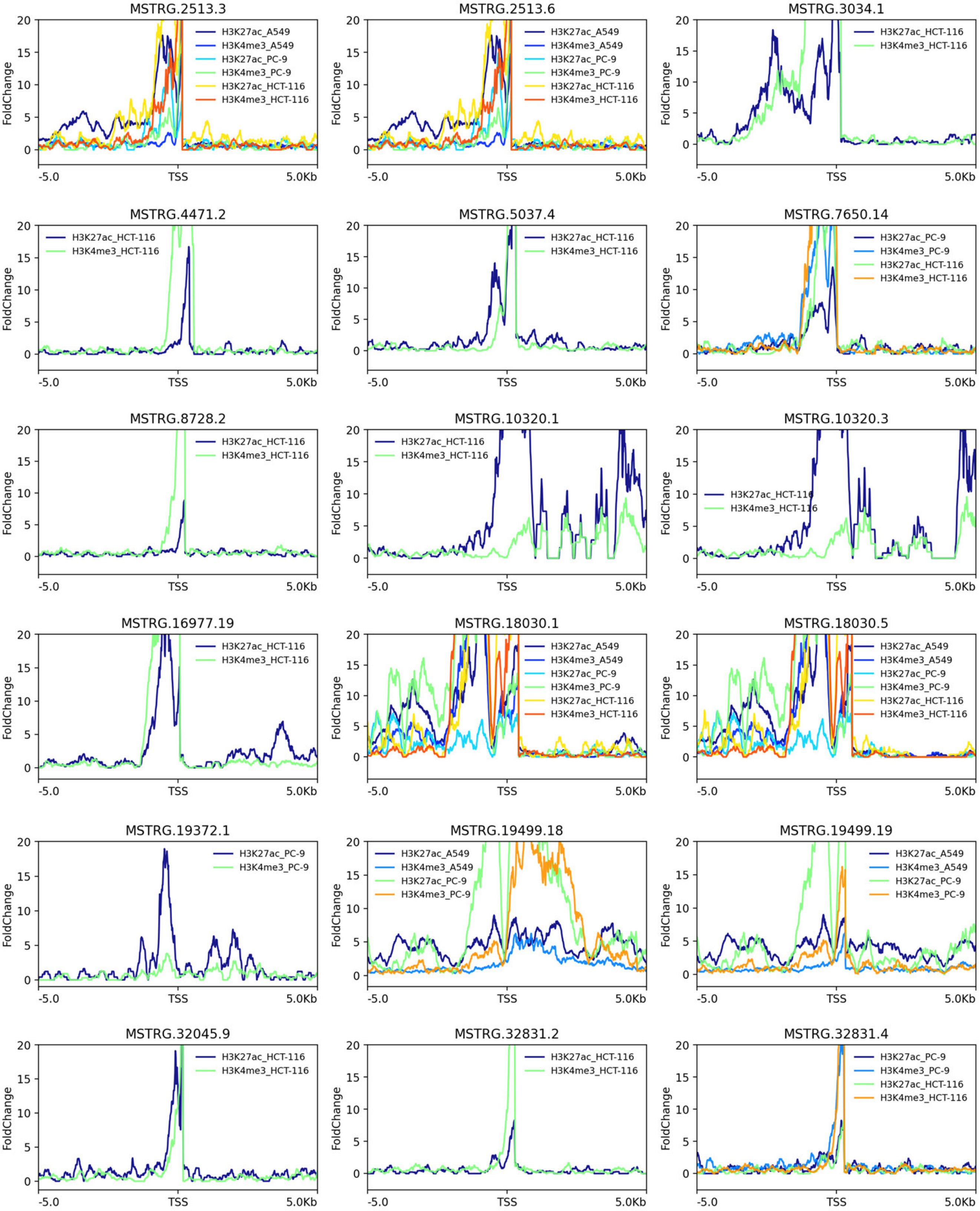


Supplementary Figure 5. H3K4me3 and H3K27ac signals near the transcription start site of the 'k' and 'j' type noncoding MSTRG transcripts in A549, HCT116, HpeG2, and PC-9 cell lines.

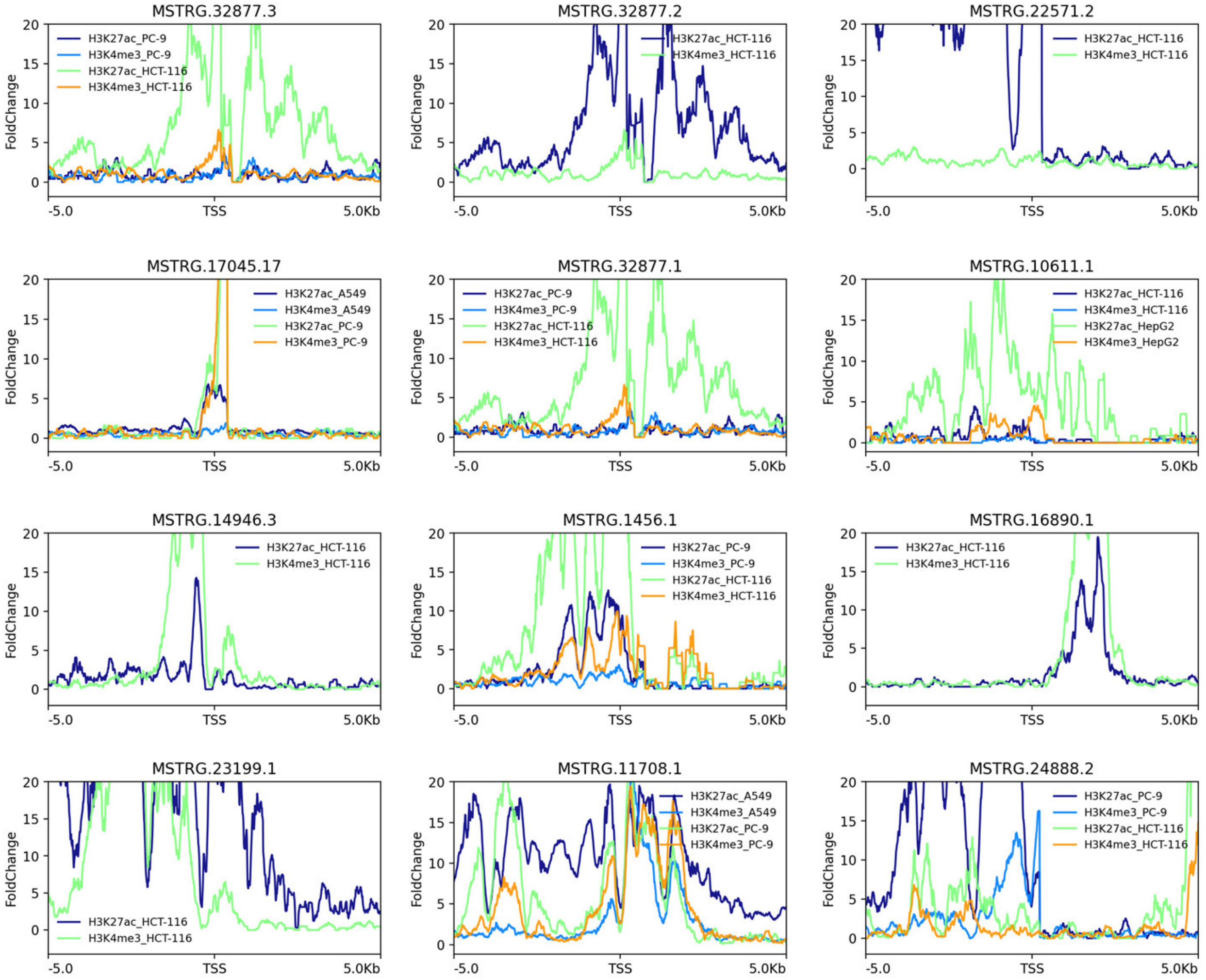


Supplementary Figure 6. H3K4me3 and H3K27ac signals near the transcription start site of the 'u' type noncoding MSTRG transcripts in A549, HCT116, HpeG2, and PC-9 cell lines.

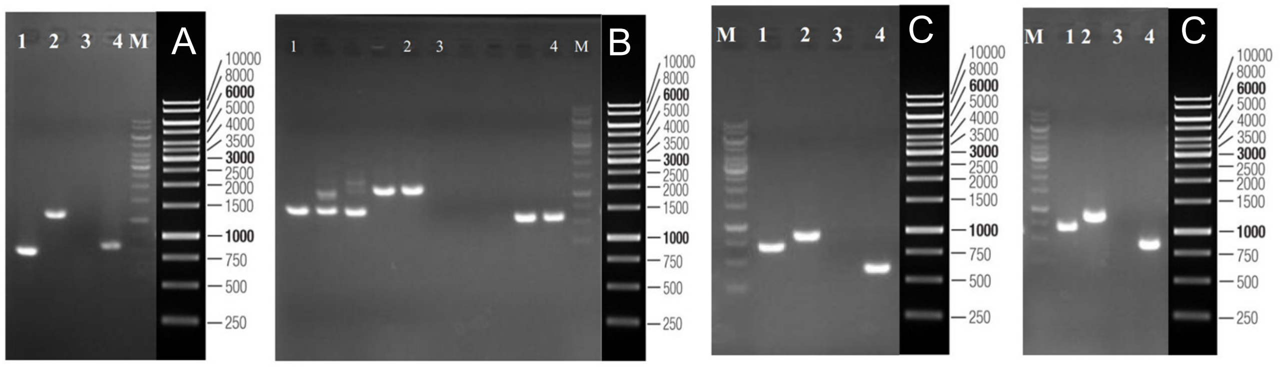


Supplementary Figure 7. DBD knockout was validated by gel electrophoresis. Shown are images of agarose gel electrophoresis. Lane 1 and lanes 2-4 indicate the KO and WT cases. Lane 2 was generated using primers far from the knockout region; lane 3 was generated using primers close to the knockout region, and the closer primer didn't generate a band. (AB) The gel electrophoresis for DBD knockout in MSTRG.2513.6 in A549. (B) The gel electrophoresis for DBD knockout in MSTRG.2513.6 in HCT116. (C) The gel electrophoresis for DBD knockout in MSTRG.4401.1 in A549. (D) The gel electrophoresis for DBD knockout in MSTRG.4401.1 in HCT116.

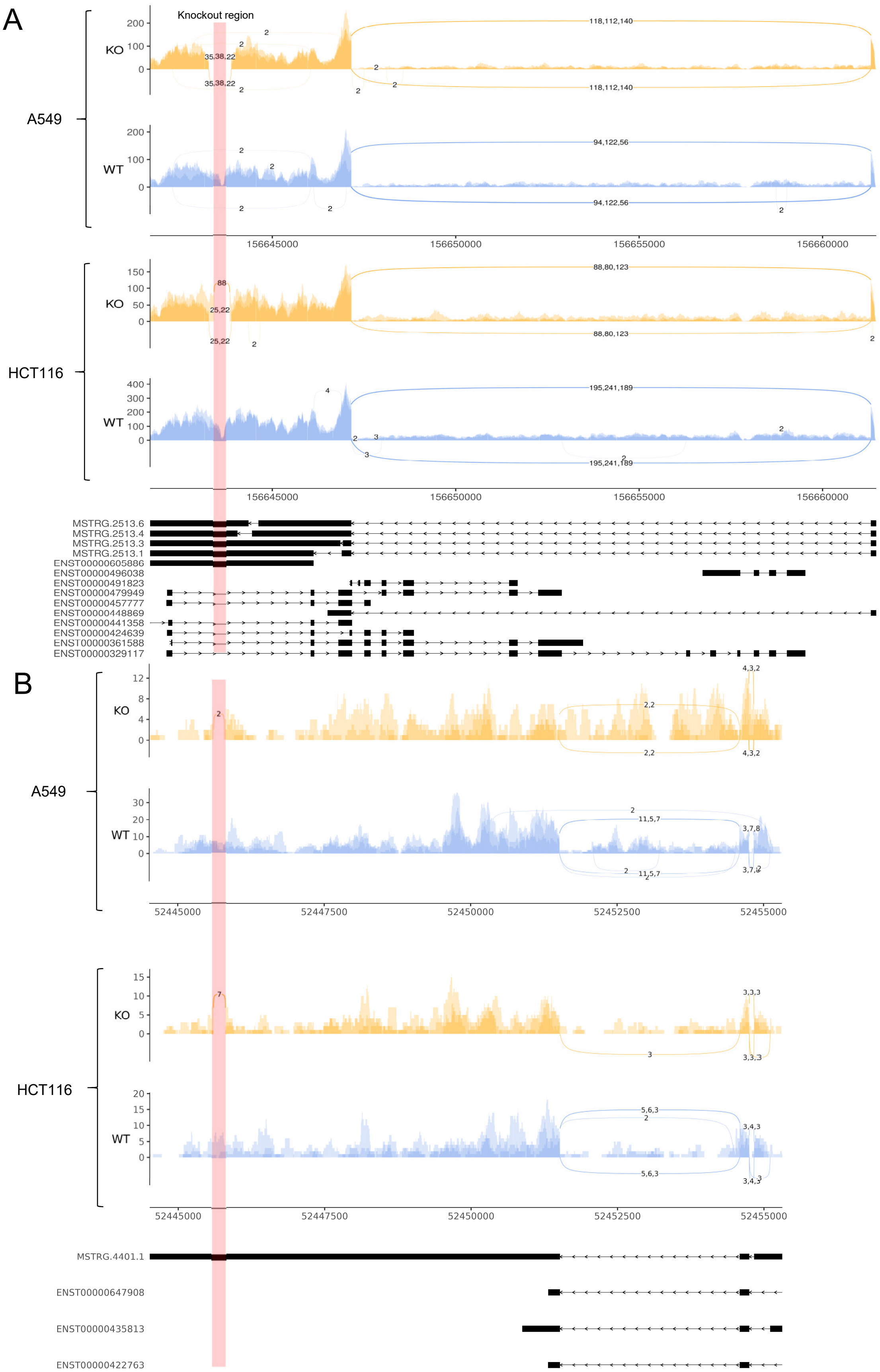


Supplementary Figure 8. The read coverage of MSTRG transcripts before and after DBD knockout (DBD regions are indicated by pink bars). (A) The reads coverage of MSTRG.2513.6. (B) The reads coverage of MSTRG.4401.1.

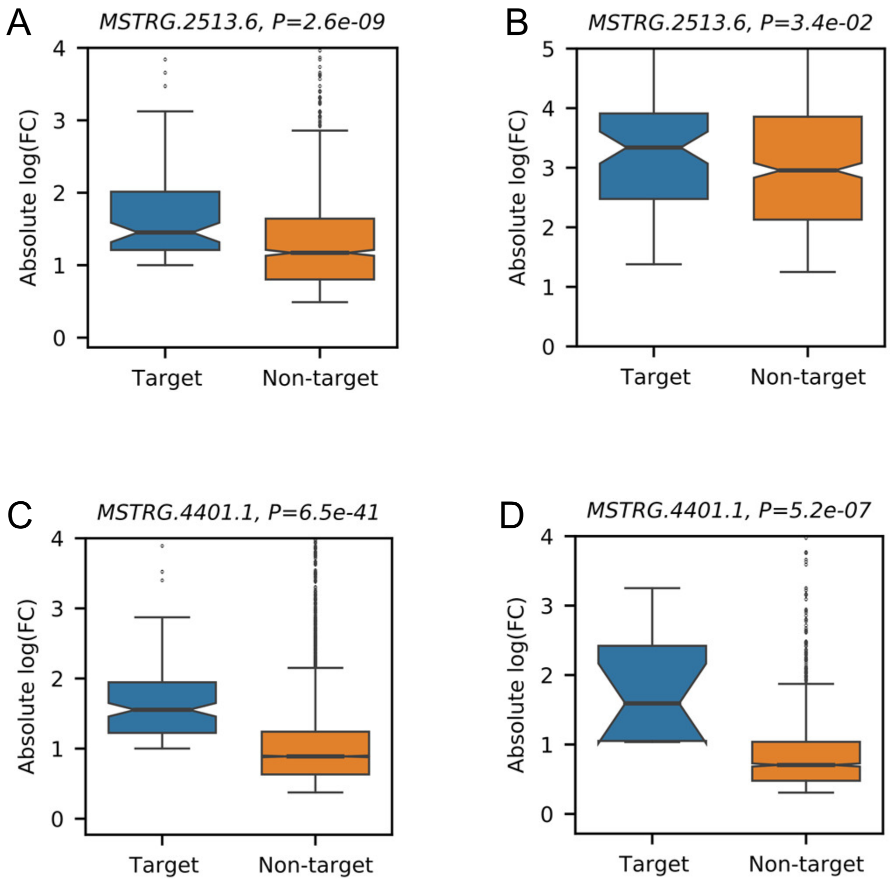


Supplementary Figure 9. The expression change of target genes was significantly larger than that of non-target genes after DBD knockout. The fold change of gene expression was computed using the edgeR package. The |fold change| distribution of target genes was compared with the |fold change| distribution of non-target genes (one-sided Mann-Whitney test). (A-D) The knockout of the DBD in MSTRG.2513.6 in A549 and HCT116. (CD) The knockout of the DBD in MSTRG.4401.1 in A549 and HCT116.

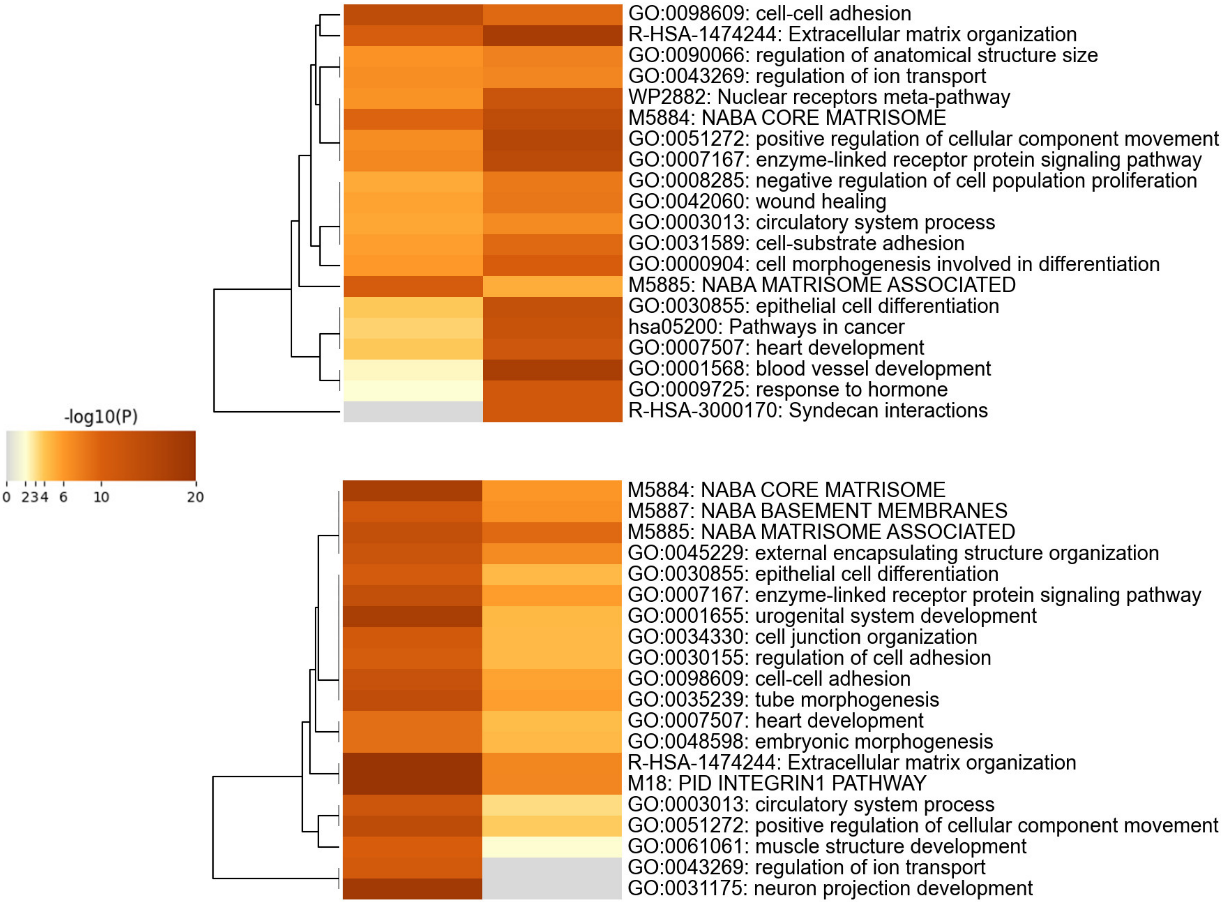


Supplementary Figure 10. The impact of DBD knockout on gene expression in cancer cell lines. The upper panel shows the cases of MSTRG.2513.6 DBD knockout in A549 and HCT116 cell lines; differentially expressed genes are commonly enriched in cell-cell adhesion and extracellular matrix organization-related gene sets and pathways. The bottom panel shows the cases of MSTRG.4404.1 DBD knockout in A549 and HCT116 cell lines; differentially expressed genes are commonly enriched in extracellular matrix-related pathways. In line with these, based on *gProfiler*, the mostly enriched gene sets and pathways were cell-matrix adhesion (GO:0007160), ECM-receptor interaction (KEGG:04512), and focal adhesion (KEGG:04510, WP306).

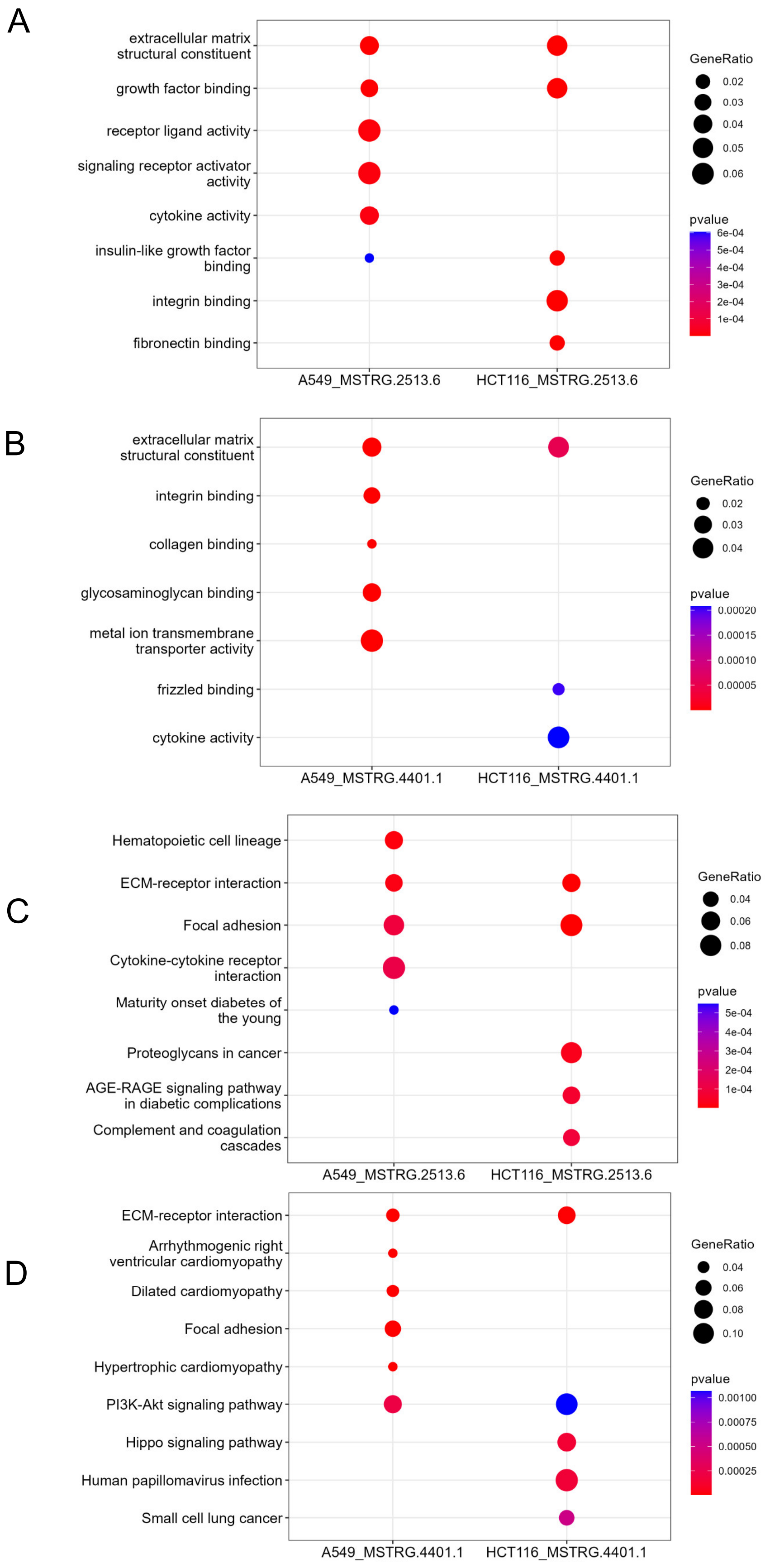


Supplementary Figure 11. Differentially expressed genes are enriched in specific KEGG pathways.

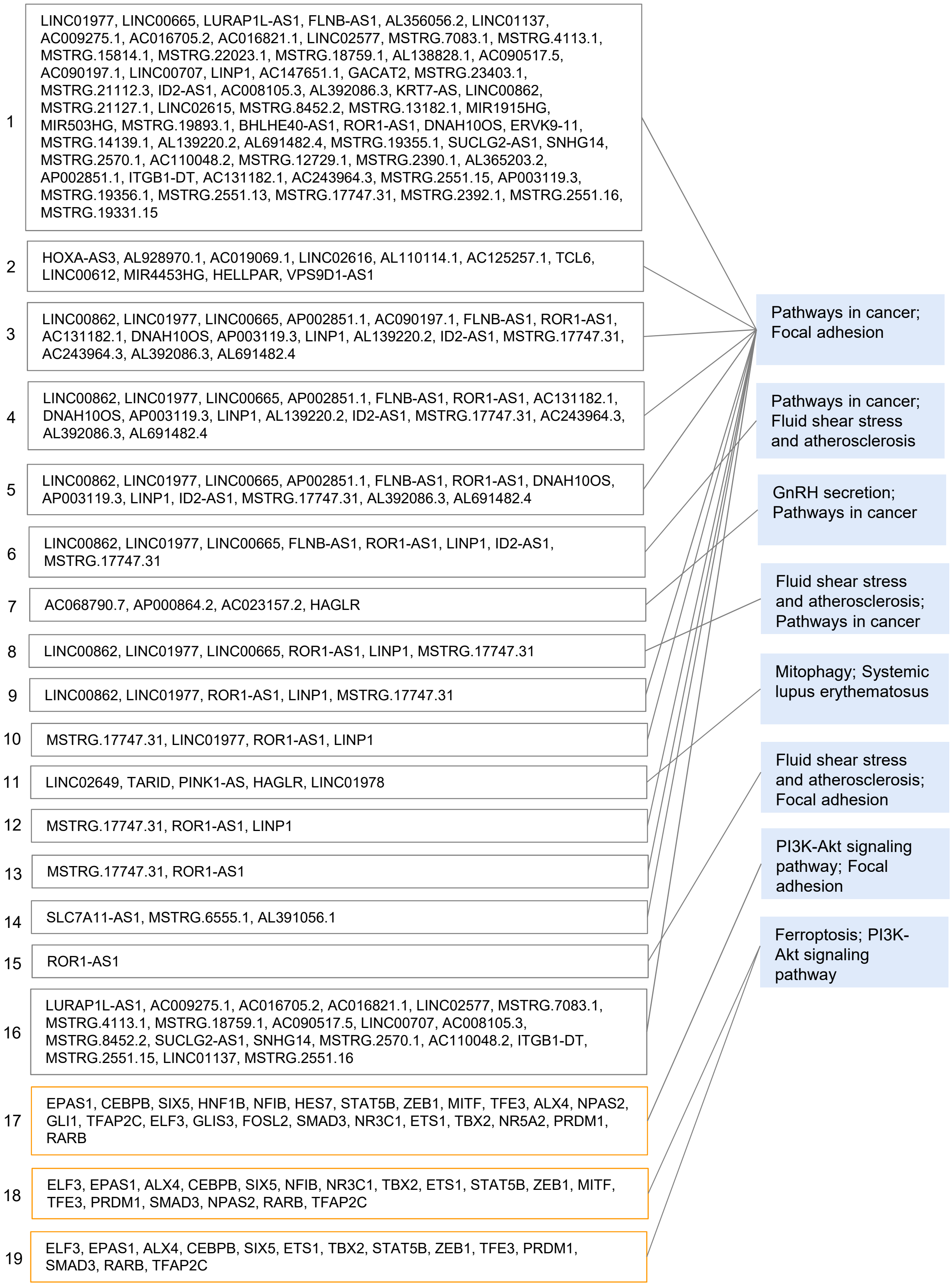


Supplementary Figure 12. The regulatory sets (lncRNA sets and TF sets), combined target modules, and the top two most enriched KEGG pathways in these modules in MSTRG.2513.6 KO HCT116. The FDR of 'Pathway in cancer' and 'Focal adhesion' ranges from 0.000161 to 0.000000 and 0.000315 to 0.000003, respectively (hypergeometric test).
